# OptiK: An Entropy-Driven Framework for Optimal k-mer Size Selection for Bacterial Genomics

**DOI:** 10.1101/2025.05.21.655412

**Authors:** AJ Gutierrez-Escobar

## Abstract

K-mer-based approaches have become fundamental (Zielezinski et al., 2017) to modern computational genomics, underpinning tools for genome assembly, metagenomic classification, variant calling, and phylogenetic analysis. Despite their ubiquity, selecting an appropriate k-mer size (k) is often made arbitrarily or heuristically, with little consideration for the underlying signal quality relative to a given dataset. Here, I introduce OptiK, a novel alignment-free tool that evaluates the information richness of k-mer encodings across a range of k values to identify the optimal k for comparative analysis. OptiK operates by constructing k-mer frequency matrices from genome collections, reducing their dimensionality via truncated singular value decomposition (SVD), and evaluating clustering structure through unsupervised metrics including the Silhouette coefficient, Calinski-Harabasz index, and Davies-Bouldin index. We validate OptiK on a curated dataset of 1044 Helicobacter pylori genomes with well-characterized population structure. OptiK robustly identifies k = 8 as the optimal k-mer size, yielding latent structures in UMAP space that align with fineSTRUCTURE-defined subpopulations without relying on prior labels or reference alignments. These results demonstrate that OptiK provides a reproducible, alignment-free strategy for optimizing k-mer resolution in bacterial comparative genomics.

## Introduction

K-mer analysis forms the computational backbone (Zielezinski et al., 2017) of many genomic methodologies, from assembly and alignment to taxonomic classification and evolutionary inference. The k-mer size, denoted k, directly shapes the resolution and interpretability of such analyses. Short k-mers may overgeneralize and conflate distinct signals, while long k-mers risk data sparsity and fragmentation. Despite this centrality, the choice of k is typically made ad hoc or copied from precedent, with little empirical justification provided.

OptiK is designed to resolve this gap. Rather than assuming a static k for all scenarios, OptiK introduces a flexible, entropy-aware framework (Arbelaitz et al., 2013) to evaluate and select the k-mer size that captures the richest, most structured signal in a dataset. Unlike methods requiring alignment, taxonomic annotation, or supervised labels, OptiK works directly with raw FASTA input and uses only unsupervised, intrinsic measures to optimize its optimization. It leverages manifold learning and clustering validation to determine which k yields the most coherent projection of sequence space.

## Methods

### Genome sequences

I downloaded the whole-genome sequences generated within the *Helicobacter pylori* Genome Project (HpGP) from GenBank BioProject PRJNA529500 (Thorell et al., 2023).

### Input Processing and k-mer Matrix Construction

OptiK begins by reading a directory of genome FASTA files. For each user-specified k-mer size (e.g., k = 3 to 8), the tool computes normalized k-mer frequency vectors for every genome. These vectors are built by sliding a window of length k across each sequence and counting the occurrences of each unique k-mer. The result is a high-dimensional feature matrix of shape (n_samples, n_kmers), where each row represents a genome and each column corresponds to a distinct k-mer. To mitigate memory overhead, OptiK dynamically prunes k-mers with zero variance across the dataset and supports sparse matrix formats. All k-mer strings are canonicalized to ensure strand symmetry by choosing the lexicographically smaller of the forward and reverse complement.

### Dimensionality Reduction via Truncated SVD

Given the high dimensionality and inherent sparsity of k-mer matrices (Alneberg et al., 2014), OptiK applies truncated singular value decomposition (SVD) to project the data into a lower-dimensional latent space. The number of retained components is set to a minimum of 50 or (n_features - 1), preserving most of the variance while enabling downstream clustering. Truncated SVD was selected over PCA because it can operate directly on sparse matrices without requiring full matrix centering.

### Optimal k is Selection

To identify the optimal k-mer size, OptiK applies a rank-based consensus approach across the three clustering validation metrics. For each tested value of *k*, the average Silhouette score, Calinski-Harabasz index, and Davies-Bouldin index (inverted for ranking, since lower is better) are computed across all cluster numbers. Each *k* is then independently assigned a rank within each metric. The overall consensus score is calculated by summing the ranks across all three metrics, and the *k* with the lowest cumulative rank is selected as optimal. In the event of tied scores, priority is given to higher Silhouette values and lower Davies-Bouldin indices, emphasizing strong intra-cluster cohesion and inter-cluster separation. This strategy ensures a balanced and interpretable selection process that avoids overreliance on any metric.

### Clustering and Metric Evaluation

Depending on user configuration, the reduced matrix is clustered using K-Means (default) or agglomerative clustering. For each k-mer size, clustering is performed independently across a fixed range of cluster numbers (e.g., n_clusters = 3 to 8). Three unsupervised clustering validation indices are calculated (Arbelaitz et al., 2013): Silhouette Score, which measures intra-cluster cohesion and inter-cluster separation; Calinski-Harabasz Index, which captures the ratio of between-cluster to within-cluster dispersion; and Davies-Bouldin Index, which penalizes cluster overlap and dispersion; lower values are better. These metrics are averaged across the clustering range for each k-mer size. The optimal k yields the highest Silhouette and Calinski-Harabasz scores and the lowest Davies-Bouldin index.

### Visualization and Output

Optionally, OptiK generates two types of visualizations per k-mer size: UMAP Projection, a two-dimensional embedding (Armstrong et al., 2021) of the reduced data colored by cluster assignment or known labels, and Hierarchical Dendrogram. When agglomerative clustering is used, the dendrogram reveals pairwise relationships between samples.

### Implementation and Reproducibility

OptiK is implemented in Python 3 and relies on standard scientific libraries: NumPy, SciPy, scikit-learn, Biopython, tqdm, and optionally UMAP. It includes robust command-line argument parsing, logging, error handling, and output checkpointing. All metric scores, cluster assignments, and visualizations are saved to the output directory. Results can be reviewed interactively or used downstream for classification, dimensionality inspection, or pipeline tuning. All results are deterministic given a random seed and reproducible via stored matrices and configuration parameters.

## Results

The overall architecture of the OptiK system is summarized in Figure 1. The workflow begins with a collection of genome sequences in FASTA format and iterates over a user-defined range of k-mer sizes. For each *k*, the system computes normalized and canonicalized k-mer frequency vectors, constructs a sparse k-mer matrix, and filters out zero-variance features. The resulting matrix is projected into a lower-dimensional latent space using truncated singular value decomposition (SVD), followed by unsupervised clustering. Cluster quality is evaluated using three internal validation metrics—Silhouette score, Calinski-Harabasz index, and Davies-Bouldin index—which are stored for each *k*. After all k values are processed, OptiK compares the metric trends and selects the k-mer size that yields the most coherent clustering structure. Optional visualizations, such as UMAP projections and dendrograms, are generated for interpretability.

**Figure 1.**
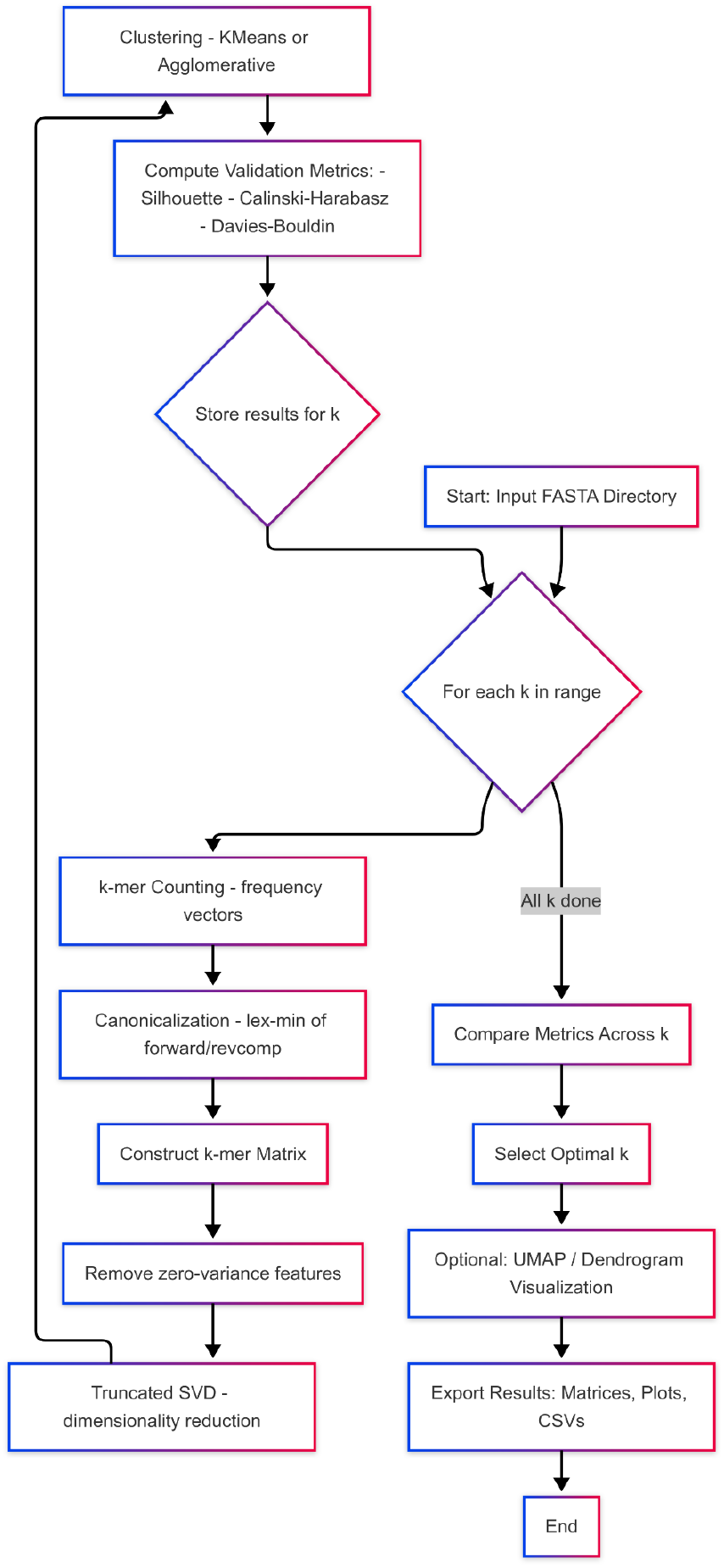
Workflow overview of the OptiK system. This flowchart illustrates the complete analytical pipeline implemented in OptiK for selecting the optimal k-mer size from a given set of genome sequences. The process begins with a directory of input FASTA files and iterates over a user-specified range of k-mer sizes. For each value of k, OptiK computes canonicalized k-mer frequency vectors, constructs a genome-by-k-mer matrix, and filters out invariant features. Dimensionality reduction is performed using truncated singular value decomposition (SVD) and unsupervised clustering using either K-Means or agglomerative methods. Clustering results are evaluated using three internal validation metrics: Silhouette score, Calinski-Harabasz index, and Davies-Bouldin index. The results for each k are stored, and the optimal k is selected based on metric performance across the tested range. The reduced feature space can be visualized via UMAP projection or hierarchical clustering. Final outputs include clustering assignments, metric plots, dimensionality-reduced matrices, and publication-ready figures.

### Optimal k-mer Discovery

I applied OptiK to the HpGP dataset, comprising 1044 publicly available genome assemblies annotated with their geographic subpopulation assignments. This dataset is a robust benchmark for biological signal recovery. To evaluate the effect of k-mer resolution on clustering quality, we applied OptiK to the HpGP dataset across k-mer sizes ranging from 3 to 8 and cluster numbers from 3 to 8. Figure 2 shows that the Silhouette score increased consistently with higher k values, peaking at *k = 8*, particularly for 7–8 clusters. The Calinski-Harabasz index followed a U-shaped trajectory, recovering at *k = 8* after a mid-range decline. In contrast, the Davies-Bouldin index decreased monotonically, indicating improved cluster compactness and separation at larger k values. Collectively, these trends converged to identify *k = 8* as the optimal k-mer size for this dataset, offering the best trade-off between resolution and structural coherence in an unsupervised setting.

**Figure 2.**
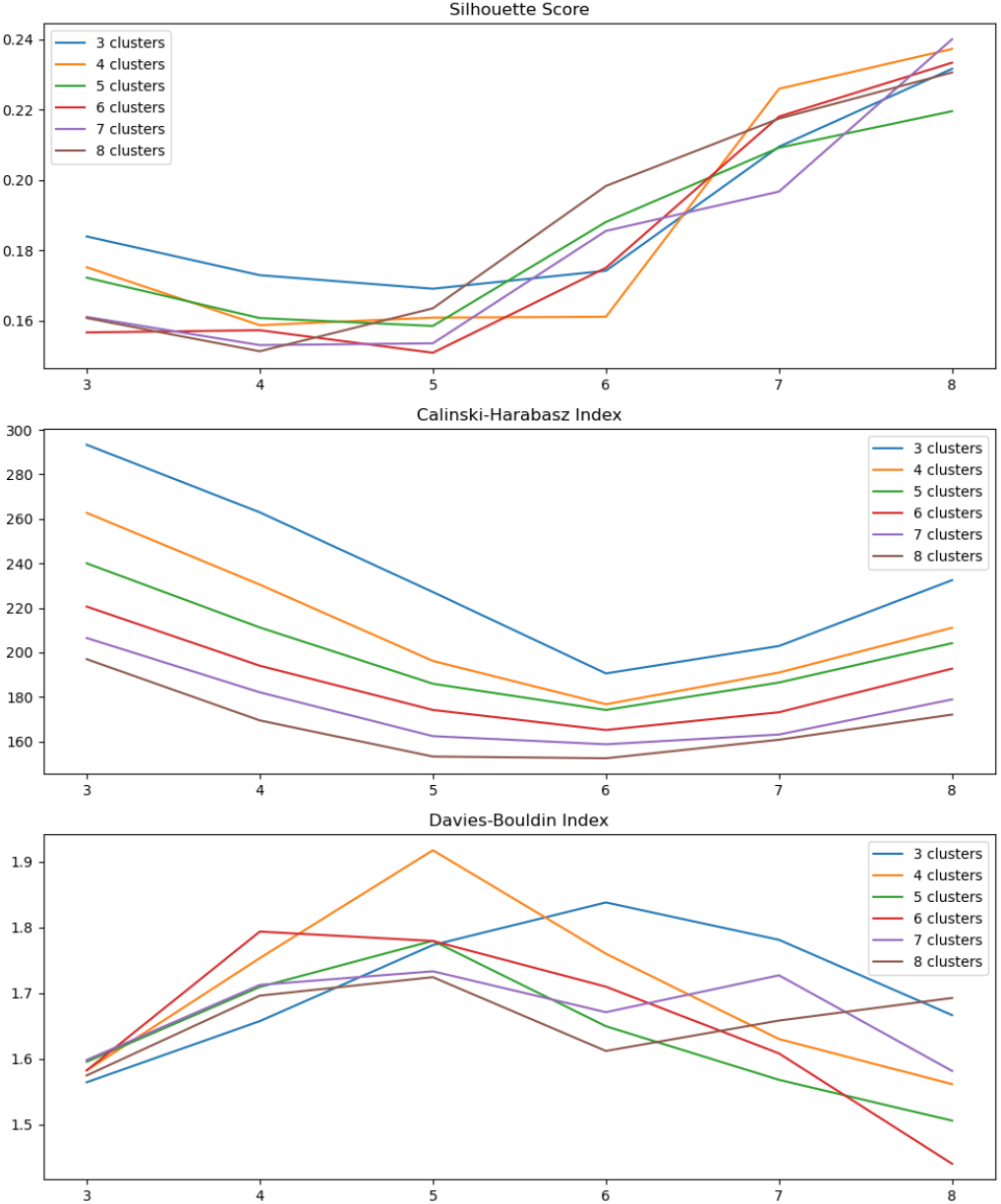
Clustering validation metrics across k-mer sizes and cluster numbers in the HpGP dataset. This panel displays three internal clustering evaluation metrics computed by OptiK across k-mer sizes ranging from *k = 3* to *k = 8*, and cluster numbers ranging from 3 to 8. (Top) Silhouette score, which quantifies the compactness and separation of clusters, increases steadily with higher k values and peaks at *k = 8*, especially with 7–8 clusters. (Middle) Calinski-Harabasz index, which measures the ratio between-cluster to within-cluster dispersion, decreases at intermediate k values but improves again toward *k = 8*, suggesting a re-emergence of global structure at higher resolution. (Bottom) The Davies-Bouldin index, where lower values indicate better clustering, consistently declines with increasing *k*, reaching a minimum at *k = 8*, particularly for 5–6 clusters. These results suggest that *k = 8* maximizes overall clustering quality and supports its selection as the optimal k-mer size for the HpGP genome collection.

To further assess the structural signal captured at each k-mer size, we generated UMAP projections of the dimensionality-reduced k-mer matrices for *k* = *3* through *k* = *8 values*, coloring each point by its unsupervised cluster label. As shown in Figure 3, increasing the k-mer size markedly improved spatial separation and cluster cohesion. At lower values (e.g., *k* = *3*–*4*), cluster boundaries are diffuse and overlapping, indicating that the encoded sequence features do not adequately distinguish genome-level differences. In contrast, projections at *k* = *7* and *k* = *8* exhibit tight, well-delineated clusters with minimal intermixing, reflecting a latent structure that aligns with known subpopulation boundaries in the HpGP dataset. These visual trends are consistent with the internal metric results and reinforce *k* = *8* as the optimal resolution for this dataset.

**Figure 3.**
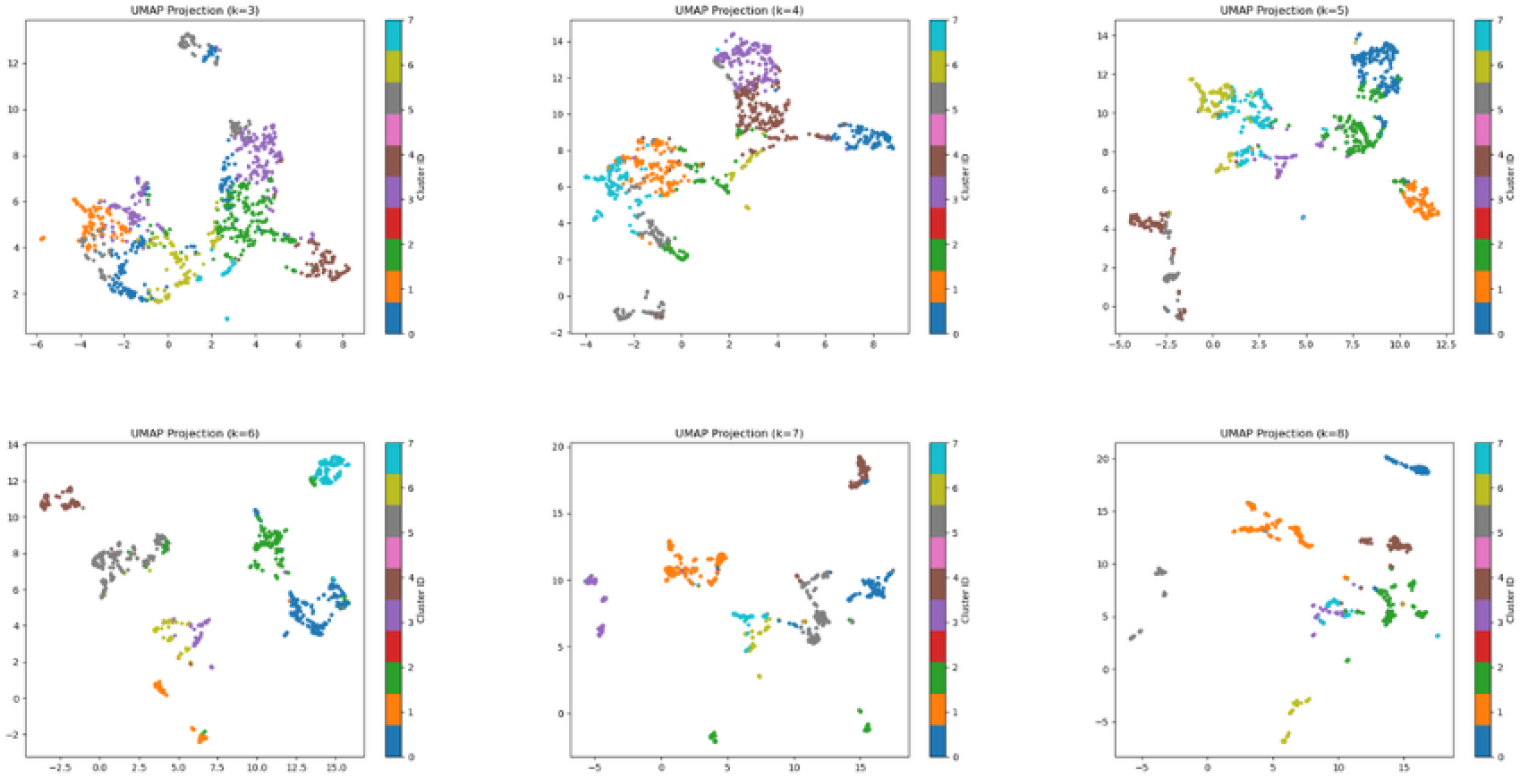
UMAP projections of dimensionality-reduced k-mer feature spaces at varying k-mer sizes (k = 3 to k = 8). Each panel shows a two-dimensional UMAP embedding of the truncated SVD-transformed k-mer frequency matrix generated by OptiK. Each point corresponds to one *Helicobacter pylori* genome (n = 1044) and is colored according to its unsupervised cluster assignment via K-Means. At lower k values (e.g., k = 3–4), clusters appear diffuse and overlapping, indicating low discriminative power. As k increases, especially at k = 7 and k = 8, clusters become more compact and well-separated, revealing meaningful latent structure in the genome collection. These projections visually corroborate the internal validation metrics and support k = 8 as the optimal resolution for this dataset.

## Discussion

The results presented here demonstrate the utility of OptiK as a principled, alignment-free tool for selecting the optimal k-mer size in genomic datasets. Despite the central role of k-mers in modern computational genomics, the problem of determining an appropriate value for *k* remains unresolved, mainly in the literature. Existing approaches typically rely on fixed heuristics or manual tuning, often without validation against structural or biological benchmarks. OptiK addresses this gap by systematically quantifying the information yield of varying k-mer sizes through unsupervised dimensionality reduction and clustering evaluation metrics.

Our application of OptiK to the Helicobacter pylori Genome Project (HpGP) dataset highlights the framework’s reliability and biological relevance. Notably, *k* = *8* emerged as the optimal k-mer resolution, supported by multiple independent metrics: the Silhouette score increased monotonically with higher *k*, the Davies-Bouldin index decreased steadily, and the Calinski-Harabasz index recovered strongly at *k* = *8*. These results were corroborated visually via UMAP projections, which revealed the emergence of well-separated and compact clusters at higher *k*, particularly between *k* = *7* and *k* = *8*. The ability of OptiK to identify this transition point—where signal clarity and structural separability converge—validates its design principles and highlights the interpretability of its output.

Although OptiK is not intended as a population structure inference tool, its outputs at the optimal *k* revealed latent structures that align with known geographic subpopulations in the HpGP dataset. This emergent correspondence suggests that entropy-informed k-mer encodings, when reduced via truncated SVD and examined through unsupervised clustering, can reflect deep genomic relationships without needing alignment or prior labels. Importantly, this observation underscores a broader implication: k-mer selection is not a neutral preprocessing choice but a decision that can significantly influence downstream biological resolution.

The UMAP projections serve a complementary function within the OptiK framework. While internal validation metrics guide the core optimization process, UMAP enables intuitive inspection of the latent space, providing researchers with a means to interpret cluster structure, assess sample relationships, and explore the extent of signal separability across values of *k*. In doing so, it bridges the gap between quantitative evaluation and visual confirmation, enhancing the overall transparency of the analysis.

This study opens several avenues for future work. First, although OptiK currently supports only frequency-based k-mer encoding, future extensions could incorporate positional or gapped k-mers and probabilistic representations informed by entropy or mutual information. Second, while the current evaluation is limited to bacterial whole genomes, the framework broadly applies to viral genomes, metagenomic bins, and transcriptomic assemblies, where k-mer dynamics vary widely in scale and composition.

While OptiK offers a robust framework for unsupervised k-mer size optimization, its current validation is limited to bacterial whole-genome datasets. It may not generalize directly to highly repetitive or fragmented genomes, such as those found in eukaryotes or metagenomic assemblies. Scalability remains a challenge at high k values or very large sample sizes due to the exponential growth of the feature space. However, dimensionality reduction via truncated SVD helps mitigate this burden. Future directions include integrating entropy-weighted k-mers, exploring scalable matrix approximations, and extending clustering options to accommodate complex genomic landscapes better, thereby enhancing OptiK’s adaptability across a broader range of comparative genomics applications.

In conclusion, OptiK provides a reproducible, scalable, and biologically coherent solution to a long-standing problem in k-mer-based genomics. By framing k-mer selection as an information optimization task and evaluating structure through unsupervised metrics, OptiK helps establish a new standard for methodological rigor in alignment-free comparative genomics. Beyond its immediate utility, OptiK lays the groundwork for broader entropy-aware sequence analytics. As genomic datasets continue to scale in volume and complexity, automated tools like OptiK that can adaptively tune foundational parameters such as k-mer size will be critical for extracting meaningful signal from noise. Its unsupervised, interpretable framework is well-positioned for integration into comparative genomics, microbial population studies, and future metagenomic pipelines.

## Availability

OptiK is freely available at https://github.com/Gutierrez-Escobar-AJ/OptiK. It includes source code, documentation, example datasets, and instructions.

## Acknowledgments

The author thanks the authors of the HpGP project.

## Notes

https://www.ncbi.nlm.nih.gov/bioproject/PRJNA529500

https://github.com/Gutierrez-Escobar-AJ/OptiK

### Competing Interest Statement

The authors have declared no competing interest.

